# Genomic surveillance of antimicrobial resistance shows cattle are a moderate source of multi-drug resistant non-typhoidal *Salmonella* in Mexico

**DOI:** 10.1101/2020.11.30.403618

**Authors:** Enrique Jesús Delgado-Suárez, Tania Palós-Guitérrez, Francisco Alejandro Ruíz-López, Cindy Fabiola Hernández Pérez, Nayarit Emérita Ballesteros-Nova, Orbelín Soberanis-Ramos, Rubén Danilo Méndez-Medina, Marc W. Allard, María Salud Rubio-Lozano

## Abstract

Multi-drug resistant (MDR) non-typhoidal *Salmonella* (NTS) is a major public health concern globally. This study reports antibiotic susceptibility testing and genotypic antimicrobial resistance (AMR) profiles of NTS isolates from bovine lymph nodes (n=48) and ground beef (n=29). Furthermore, we compared genotypic AMR data of our isolates with those of publicly available NTS genomes from Mexico (n=1637). The probability of finding MDR isolates was higher in ground beef as compared to lymph nodes: χ^2^=12.0, P=0.0005. The most common resistant phenotypes involved tetracycline (40.3%), carbenicillin (26.0%), amoxicillin-clavulanic acid (20.8%), chloramphenicol (19.5%) and trimethoprim-sulfamethoxazole (16.9%), while over 55% of the isolates showed decreased susceptibility to ciprofloxacin and 26% were MDR. Occurrence of MDR isolates was strongly associated with NTS serovar (χ^2^=24.5, P<0.0001), with Typhimurium accounting for 40% of MDR strains. Most of these (9/10), carried *Salmonella* genomic island 1, which harbors multiple AMR genes (*aadA2, blaCARB-2, floR, sul1, tetG*) that confer a penta-resistant phenotype. Moreover, 100% of MDR isolates had mutations in the *ramR* gene. Among public NTS isolates from foods and clinical cases in Mexico to date, those from cattle had the highest proportion of MDR genotypes. Our results suggest attaining significant improvements in AMR meat safety may require the identification and removal (or treatment) of product harboring MDR NTS, instead of screening for *Salmonella* spp. or for isolates showing resistance to individual antibiotics. In that sense, massive integration of whole genome sequencing (WGS) technologies in AMR surveillance provides the shortest path to accomplish these goals.

## Introduction

For decades, experimental data has supported concerns that antimicrobials use (AMU) in animal production is a key factor contributing to the ever-growing crisis of bacterial antimicrobial resistance (AMR). However, there is limited evidence linking food consumption with AMR emergence in humans [1, 2]. Moreover, an increasing number of studies have found that AMU in food animals has a limited impact on the AMR profile of foodborne pathogens [3–6]. While this may be true in developed countries, the situation is likely different in low and middle-income countries (LMIC) where controls of AMU in veterinary practice and human health are less rigorous. Therefore, in the context of an intense trade of foods between countries differing in AMR food safety, it is vital to identify relevant sources of AMR pathogens along the food chain. This would help to prevent their dissemination, as well as human exposure to multi-drug resistant (MDR) bacteria, which is a global public health issue.

In Mexico, source attribution of foodborne illnesses is in an early stage. Nonetheless, official epidemiological surveillance reports high incidence rates for infections commonly transmitted through foods. That of non-typhoidal salmonellosis has been above 60 cases per 100 thousand inhabitants in the last 5 years [7]. In addition, recent studies conducted in Mexico show non-typhoidal *Salmonella* (NTS) contamination is unusually high (15 to nearly 100% positive samples) in bovine lymph nodes, beef carcasses and ground beef, with isolates exhibiting rates of resistance that vary from 30 to nearly 98%, across several important antimicrobial classes (i. e. penicillins, aminoglycosides, cephalosporins, quinolones) [8, 9]. This evidence suggests beef is likely to play a role as a relevant reservoir of foodborne MDR salmonellosis in Mexico. Especially, if considering it is often involved in salmonellosis outbreaks in countries with lower NTS contamination rates [10, 11].

The increasing availability of whole genome sequencing (WGS) technologies have helped to improve genomic surveillance as it provides a high-resolution method for the characterization of an organism features. Regarding AMR in NTS, however, most research from LMIC deal with phenotypes [12–16]. Although these studies raise concerns, they do not provide insights into the genetic basis of AMR, its evolution or dissemination within bacterial populations. Such information is crucial to devise new strategies to contain the dissemination of AMR pathogens. However, it can best be obtained by addressing phenotypic and genomic profiling of AMR simultaneously, an area with a limited number of studies in Mexico and other countries [17].

In the present investigation, we conducted antibiotic susceptibility testing and WGS of 77 NTS isolates collected in the course of a previous research project involving bovine lymph nodes (n=800) and ground beef (n=745) across a two-year sampling period [18]. Assembled genomes were used to predict the AMR genomic profiling of NTS isolates, and this was further compared to their corresponding AMR phenotypes. We also conducted comparative genomics of AMR genotypes of publicly available NTS isolates collected in Mexico from bovines, vegetables, seafood, aquatic environments, human stools and clinical cases. Consequently, we managed to thoroughly characterize the role of cattle as a reservoir of NTS harboring AMR genes of human clinical significance. Moreover, we identified the dominant genetic determinants that sustain resistance against specific antibiotic classes, as well as MDR phenotypes.

## Materials and Methods

Animal Care and Use Committee approval was not obtained for this study since live animals were not directly involved in the experiment.

### *Salmonella* isolates

We used 77 strains from a previous study conducted by our research group [18]. These were isolated from bovine lymph nodes (LN, n= 800) and ground beef (GB, n=745) collected between April 2017 and December 2018. Each isolate was obtained from a different sample across the two-year sampling period.

#### Antibiotic Susceptibility Testing (AST)

The phenotypic AMR profile of NTS isolates was determined with a panel of 13 antibiotics included in the WHO list of critically important and highly important antimicrobials [19]. We used the disk diffusion method [20], with the Bencton Dickinson AST test disks and concentrations reported in Table 1.

**Table 1.** Antimicrobial agents tested, their concentrations and zone diameter clinical break points used in the AST

| Antimicrobials | C, µg<sup>2</sup> | Zone diameter breakpoint<sup>1</sup>, mm |  |
| --- | --- | --- | --- |
|  |  | R | I |
| Amikacin (AMK) | 30 | ≤14 | 15-16 |
| Gentamicin (GEN) | 10 | ≤12 | 13-14 |
| Carbenicillin (CB) | 100 | ≤19 | 20-22 |
| Amoxicillin/clavulanic acid (AMC) | 20/10 | ≤13 | 14-17 |
| Cefotaxime (CTX) | 30 | ≤22 | 23-25 |
| Ceftriaxone (CRO) | 30 | ≤19 | 20-22 |
| Cefepime (FEP) | 30 | ≤18 | 19-24 |
| Imipenem (IPM) | 10 | ≤19 | 20-22 |
| Ertapenem (ETP) | 10 | ≤18 | 19-21 |
| Meropenem (MEM) | 15 | ≤19 | 20-22 |
| Chloramphenicol (CHL) | 30 | ≤12 | 13-17 |
| Trimethoprim-sulfamethoxazole (SXT) | 1.25/23.75 | ≤10 | 11-15 |
| Ciprofloxacin (CIP) | 5 | ≤20 | 21-30 |
| Tetracycline (TET) | 30 | ≤11 | 12-14 |

<sup>1</sup>Criteria to consider isolates as clinically resistant (R) or intermediate (I) [21]. For meropenem, we also considered epidemiological cutoff values (ECOFF) set by EUCAST to classify strains as wild-type (>27 mm) or non-wild-type (≤27 mm) [22].
<sup>2</sup>Antimicrobial's disk concentration.

Isolates were classified as susceptible, intermediate or resistant, according to the Clinical Laboratory Standards Institute (CLSI) guidelines [21]. Strains of *Escherichia coli* ATCC 8739, *Enterococcus fecalis* ATCC 29212, *Staphylococcus aureus* ATCC 25923, and *Pseudomonas aeruginosa* ATCC 9027 were used as quality control organisms. Isolates showing intermediate and resistance phenotypes were considered as non-susceptible in this study. Likewise, isolates showing resistance to ?3 antimicrobial classes were classified as MDR [23]. The detailed AST protocol is available at protocols.io (doi: dx.doi.org/10.17504/protocols.io.bpypmpvn).

### Whole-genome sequencing and genome assembly

Genomic DNA (gDNA) was extracted from fresh colonies grown overnight at 37° C in tryptic soy broth. For that purpose, we used the Roche PCR High Purity Template Preparation Kit (Roche México, Mexico City, Mexico), following manufacturer’s instructions. Subsequently, gDNA was quantified using a Qubit 3.0 Fluorometer (Thermo Fisher Scientific México, Mexico City, Mexico). Afterwards, sequencing libraries were prepared from 1 ng gDNA using the Nextera XT Library Preparation Kit v.3 (Illumina) and sequenced on the Illumina NextSeq system (paired end 2 x 150 bp insert size). Raw sequences are publicly available at the National Center for Biotechnology Information (NCBI). The accession numbers and metadata are listed in supplementary S1 Table.

The quality of raw reads was first assessed with FastQC [24] and we used Trimmomatic [25] to filter poor-quality reads and Illumina adaptors. Trimmed sequences were analyzed again with FastQC to ensure only high-quality reads (i. e. Q≥30) were used for *de novo* genome assembly. Finally, genome assembly was conducted in the PATRIC web server [26] using the SPAdes algorithm [27].

### Genotypic AMR profiling and comparative genomics

AMR genes and point mutations associated with AMR were predicted with the aid of AMRFinderPlus program version 3.8 using assembled genomes [28]. The study also aimed to compare the genetic AMR profile of our isolates in the context of NTS population circulating in the country. For that purpose, we identified all *Salmonella* isolates from Mexico that were publicly available at NCBI as of September 21, 2020 (n=2400). Among these, we selected groups of isolates of known isolation source: bovines (n=114, including the 77 from this study), vegetables (n=1,064), seafood (n=129), aquatic environment (n=385), and human stools/clinical cases (n=22). The human isolates group will be referred to as “human clinical cases” from now on. The full list of accessions and metadata of these isolates is provided in supplementary S2 Table. AMR genotypes of these isolates were collected from the NCBI Pathogen Detection web site (https://www.ncbi.nlm.nih.gov/pathogens), which are generated by the AMRFinderPlus database and program.

Most serovar Typhimirum isolates (8/10) exhibited a penta-resistant phenotype similar to that reported for the Typhimurium DT104 strain, which is sustained by *Salmonella* genomic island 1 (SGI1) [29]. Hence, we conducted a BLAST atlas analysis to determine if the MDR profile of these isolates was associated with this genomic feature. For that purpose, we used assembled genomes at the GView web server [30], configured as follows: expect e-value cutoff=0.001, genetic code=bacterial and plant plastid, alignment length cutoff=50, percent identity cutoff=70 and tblastx as the BLAST program. The SGI1 reference sequence (AF261825.2) was collected from the Pathogenicity Island Database [31]. Furthermore, considering the epidemiological importance of this serovar, we also analyzed the whole set of Typhimurium isolates from Mexico deposited at NCBI (n=38, refer to S3 Table for accession numbers and AMR genotypes of this group of isolates). This analysis was performed in order to estimate how common MDR profiles are in strains of this serovar circulating in Mexico.

### Plasmid profiling

Plasmids are known as strong contributors to AMR dissemination. Hence, we used PlasmidFinder 2.1 [32] to predict the isolates plasmid profile using assembled genomes and a threshold identity of 95%. When plasmid replicons were detected, we collected each plasmid’s reference sequence from NCBI and mapped it against assembled genomes. If most plasmid sequences (>70%) were represented in a genome and the genomic context of genes matched that of the plasmid, these isolates were proposed to carry the predicted plasmid.

### Data analysis

We used Chi-square tests and odds ratio calculations to determine if there was an association between AMR profiles (both phenotypic and genotypic) and NTS serovars and/or isolation sources. These analyses were performed both with our experimental isolates, as well as with the additional public NTS genomes from Mexico included in the study.

We also conducted a correlation analysis between phenotypic and genotypic AMR results. Moreover, the genomic AMR profile of public NTS from Mexico was analyzed with the aid of a heatmap. For that purpose, we first determine the number of isolates from each source harboring specific AMR genes. Using these figures, we calculated the proportion of isolates from each source having the feature and used these data to generate a heatmap with the Heatmapper software [33].

## Results

### Phenotypic and genotypic antimicrobial resistance profiles

Approximately three quarters of the isolates were non-susceptible to at least one antibiotic class, while 26% were susceptible to all the tested antibiotics (Fig 1). The most common resistant phenotypes included tetracycline (40.3%), carbenicillin (26.0%), amoxicillin-clavulanic acid (20.8%), chloramphenicol (19.5%) and trimethoprim-sulfamethoxazole (16.9%). Resistance to cephalosporins and aminoglycosides was less frequent (1.3-7.8% across these antibiotic classes), while for carbapenems only intermediate resistance was observed at a frequency of 1.3 and 9.1% for ertapenem and imipenem, respectively. On the other hand, although only one isolate resisted ciprofloxacin, 54.5% of the strains showed intermediate resistance to this drug.

**Fig 1.**
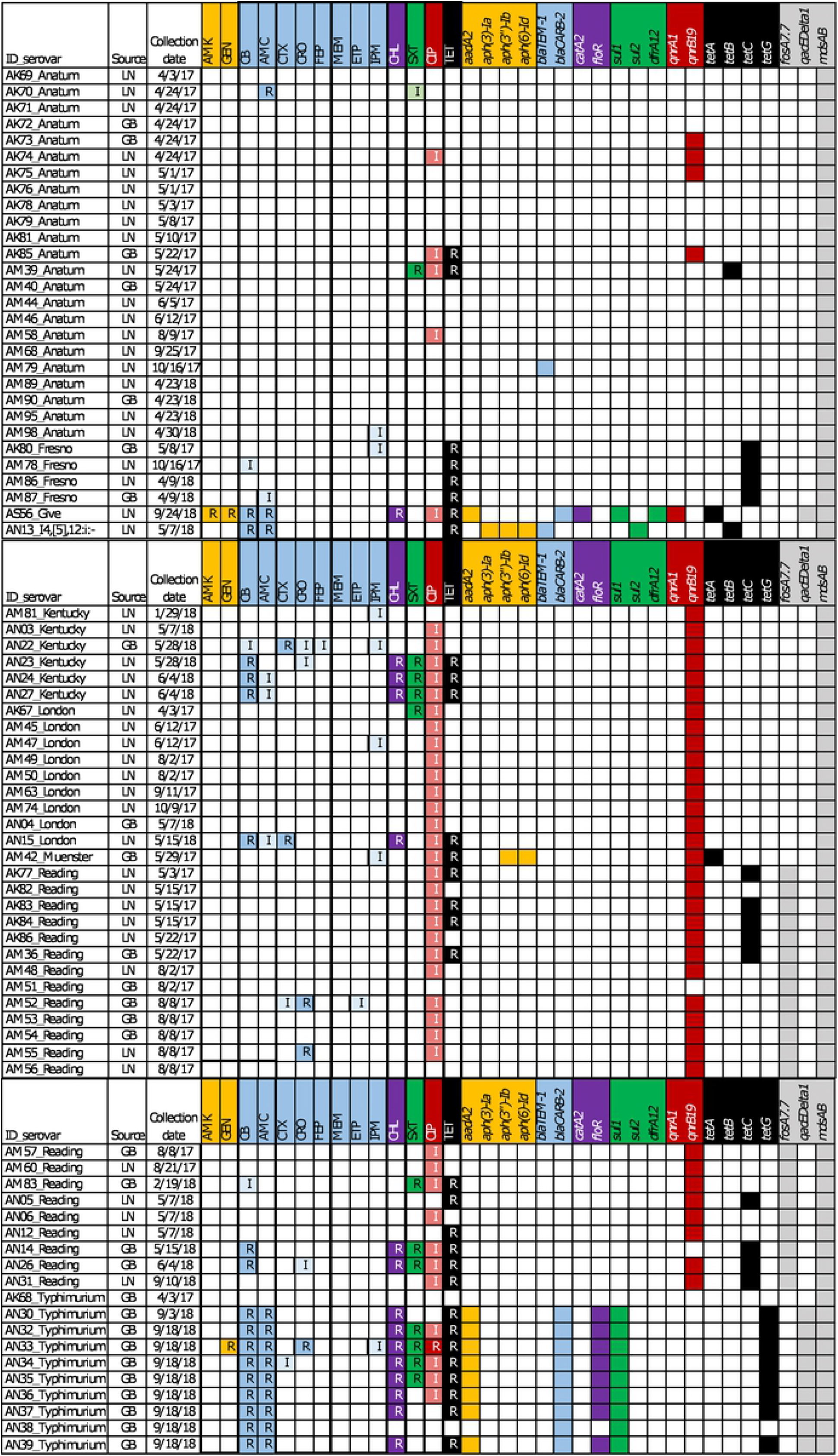
AMR profile of 77 *Salmonella* isolates from lymph nodes (LN) and ground beef (GB). Cells are color coded according to antibiotic class: orange, aminoglycosides; blue, beta-lactams; purple, phenicols; green, sulfonamides; red, fluoroquinolones; black, tetracyclines. Resistant strains are indicated with “R” and intermediate with “I”. For genotypes, cells with the antibiotic class color indicate presence of the gene. Genes in gray are not associated with any antibiotic class included in the AST panel. Antibiotic abbreviations are as follows: amikacin (AMK), gentamycin (GEN), carbenicillin (CB), amoxicillin-clavulanic acid (AMC), cefotaxime (CTX), ceftriaxone (CRO), cefepime (FEP), meropenem (MEM), ertapenem (ETP), imipenem (IMP), chloramphenicol (CHL), trimethoprim-sulfamethoxazole (SXT), ciprofloxacin (CIP), tetracycline (TET).

The rate of MDR strains was 26%, with the most common MDR profiles involving penicillins, chloramphenicol, trimethoprim-sulfamethoxazole, ciprofloxacin and tetracycline. Strikingly, there was one serovar Typhimurium isolate that resisted all antibiotic classes but carbapenems. In fact, the probability of isolates having MDR profiles was significantly higher in strains of serovar Typhimurium as compared to other serovars (OR=45.8, 95CI 5.3-399.2, P<0.0001). Likewise, there was a higher probability of finding MDR strains in ground beef as compared to lymph nodes (OR=6.5 95CI 2.1-20.1, P=0.0005).

The WGS-based *in-silico* prediction of non-susceptible phenotypes was generally good, as shown by the strong correlation between AMR genotypes and phenotypes (Fig 2).

**Fig 2.**
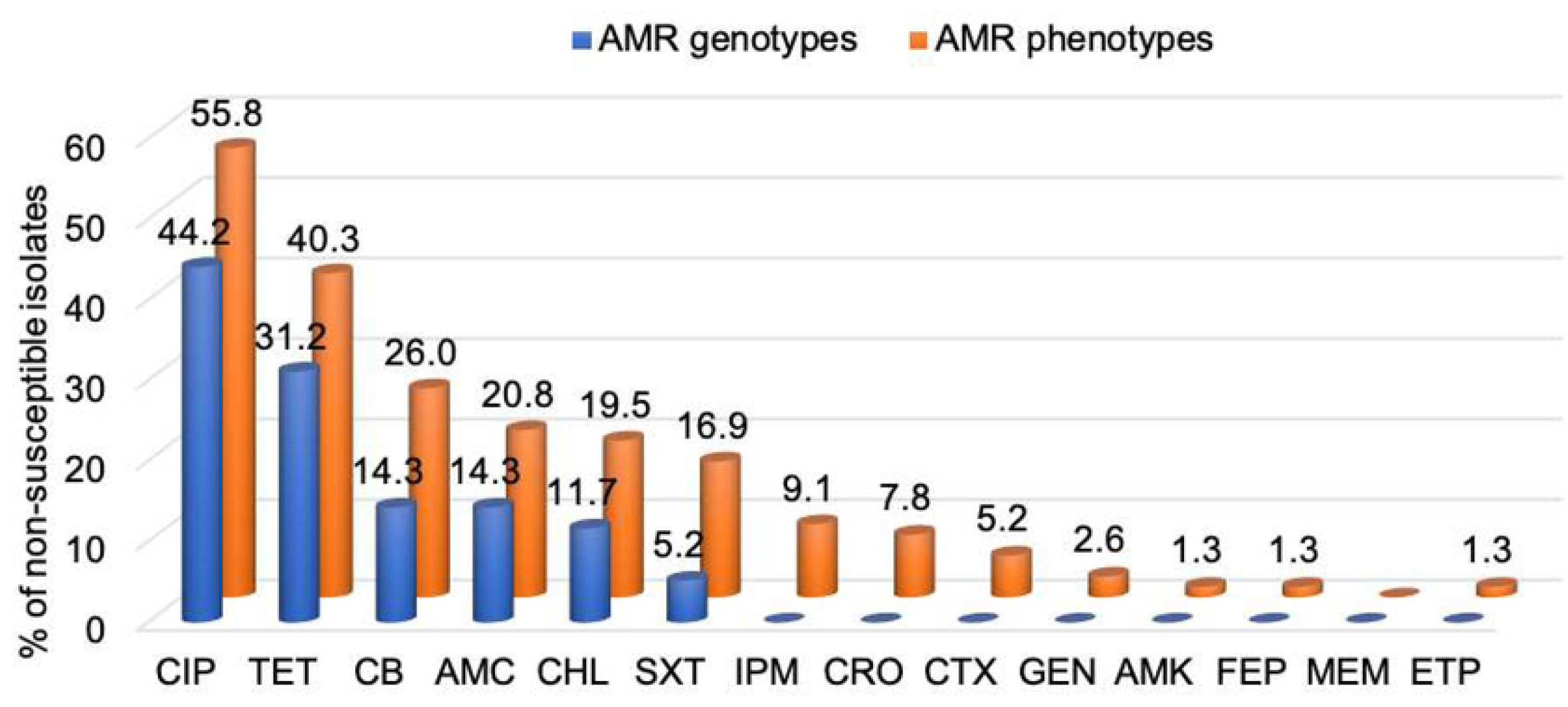
Overall correlation of phenotypic resistance with the presence of antimicrobial resistance genes for the 77 *Salmonella* isolates (r=0.979, P<0.0001). Antibiotic abbreviations: ciprofloxacin (CIP), tetracycline (TET), carbenicillin (CB), amoxicillin-clavulanic acid (AMC), chloramphenicol (CHL), trimethoprim-sulfamethoxazole (SXT), imipenem (IMP), ceftriaxone (CRO), cefotaxime (CTX), gentamycin (GEN), amikacin (AMK), cefepime (FEP), meropenem (MEM), ertapenem (ETP).

Genomic AMR profiling also identified the presence of several additional resistance genes that are not associated with any specific antimicrobial included in the AST panel. Particularly, those encoding resistance to fosfomycin (*fosA7.7*), quaternary ammonium compounds (*qacEDelta1*), as well as components of the multidrug and metal RND-type efflux complex (*mdsAB*).

Assembled genomes were also analyzed for point mutations associated with resistance. However, the occurrence of mutations was mostly inconsistent with the observed phenotypes (Table 2).

**Table 2.** Relative frequency of chromosomal point mutations detected in the studied *Salmonella* isolates (n=77) and their associated resistance phenotypes^1^

| Gene | Detected point mutations | Relative frequency, % | Resistance phenotype |
| --- | --- | --- | --- |
| *gyrA* | A67P, D72G, V73I, G81C, G81D,<br>G81H, G81S, D82G, D82N,<br>S83A, S83F, S83Y, D87G, D87K,<br>D87N, D87Y, S97P, L98V,<br>A119E, A119S, A139A | 100.0 | Quinolones |
| *gyrB* | Y421C, R438L, S464T, S464Y,<br>E466D | 100.0 | Quinolones |
| *parC* | T66I, G78D, S80I, S80R, E84G,<br>E84K, W106G | 1.3 | Quinolones |
| *parE* | M438I, E454G, S458P, V461G,<br>H462Y, A499T, V514G, V521F | 100.0 | Quinolones |
| *pmrA* | G15R, G53E, G53R, R81C,<br>R81H | 98.7 | Colistin |
| *pmrB* | L14F, L22P, S29R, T92A, P94Q,<br>E121A, S124P, N130Y, T147P,<br>R155P, T156M, T156P, V161G,<br>V161L, V161M, E166K, M186I,<br>G206R, G206W, S305R | 98.7 | Colistin |
| *acrB* | R717L, R717Q | 88.3 | Azithromycin |
| *soxR* | R20H, G121D | 100.0 | Penicillins, phenicols,<br>quinolones, rifampin,<br>tetracyclines |
| *soxS* | E52K | 100.0 | Penicillins, phenicols,<br>quinolones, rifampin,<br>tetracyclines |
| *ramR* | T18P, G25A, R46P, T50P, Y59H,<br>M84I, E160D | 68.8 | Penicillins, phenicols,<br>quinolones, rifampin,<br>tetracyclines |

<sup>1</sup>As reported by *in silico* analysis of assembled genomes with AMRFinderPlus 3.8[28]

For instance, all isolates carried multiple mutations in the quinolone resistance-determining region (QRDR), *gyrAB* and *parE* genes, regardless of being susceptible or not to ciprofloxacin. Likewise, 100% isolates had mutations in *soxRS* genes, which are known to confer MDR profiles [34]. Still, only 26% of isolates were classified as MDR. Conversely, mutations in *ramR* were strongly associated with the occurrence of MDR strains (χ^2^=17.7, P<0.0001).

Genomic analysis also revealed additional widespread mutations. Those of *pmrAB* genes, which are associated with colistin resistance, were present in all isolates. Likewise, mutations in the *acrB* gene, which confer resistance to azithromycin, were detected in 88.3% of the strains. These two drugs, however, were not included in the AST panel. Below, a detailed description of the relationship between AMR phenotypes and genotypes will be presented for each of the tested antibiotic classes, as well as for MDR strains.

### Aminoglycoside resistance

Aminoglycoside resistance was supported by the presence of genes encoding enzymatic inactivation mechanisms, such as phosphorylation (several *aph* alleles) and adenylation (*aadA2* gene). Comparative analysis with public *Salmonella* genomes revealed a strong association between aminoglycoside resistant and the source of the isolates (χ^2^=89.2, P<0.0001).

Clinical strains had the highest proportion of aminoglycoside resistance genotypes (Fig 3) and were more likely to carry AMR genes against this antibiotic class as compared to those from food related sources (OR=7.7, 95CI 3.2-18.3). Among non-clinical samples, bovines were the most significant contributors of aminoglycoside resistant NTS (OR=5.0, 95CI 3.1-7.9). In general, however, the resistance profile was similar across sources, with a predominance of *aadA* and *aph* genes over the *acc* alleles.

**Fig 3.**
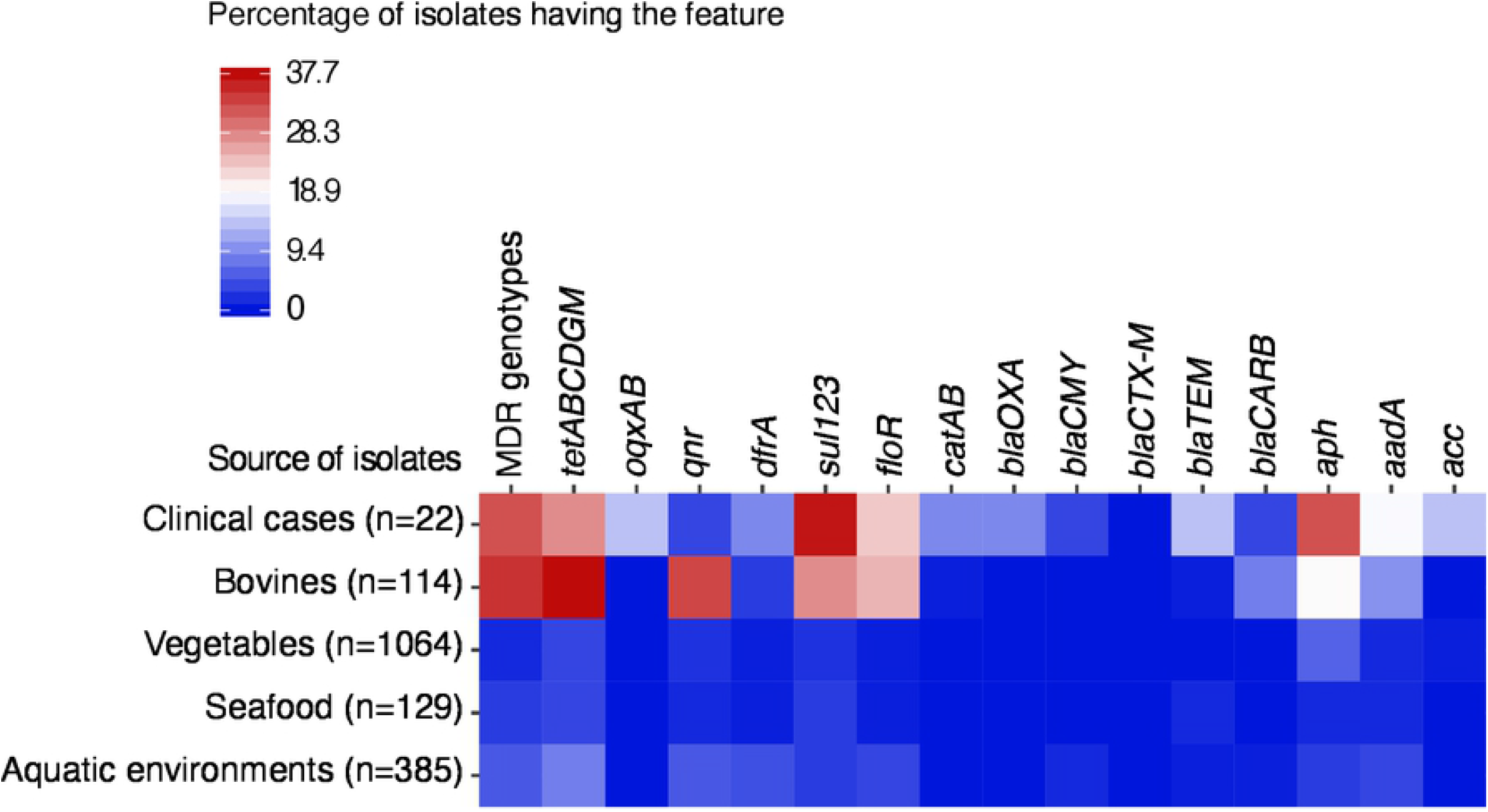
Comparative genomic AMR profile of public NTS isolated from human clinical cases, foods, and food related sources in Mexico. Refer to S2 Table for accessions and metadata of isolates included in this analysis.

### Beta-lactam resistance

Experimental isolates harbored only the Ambler’s class A beta-lactamase-encoding genes *blaCARB-2* and *blaTEM-1*. These enzymes are known to confer resistance to all penicillins, as well as first, second and third generation cephalosporins [35]. The *blaCARB-2* gene was the most abundant among isolates showing resistance to penicillins, especially in those of serovar Typhimurium (9/10). One isolate of serovar Anatum and one of monophasic (I4,[5],12:i:-) Typhimurium also carried *blaTEM-1*, which encode another extended-spectrum beta-lactamase (ESBL) that hydrolyzes penicillins and first-generation cephalosporins [35]. There were 11 non-susceptible isolates that did not carry any of the known ESBL-encoding genes. Strikingly, however, 10 out of these isolates had *ramR* mutations, which are associated with resistance towards several antibiotic classes, including penicillins.

Despite eight isolates were non-susceptible to third generation cephalosporins (3GC) and one to both 3GC and fourth generation cephalosporins (4GC), none carried ESBL-encoding genes associated with these phenotypes. As observed with penicillins, mutations in *ramR* appeared to be associated with non-susceptibility to cephalosporins since 100% of non-susceptible isolates to 3GC/4GC had these mutations. Non-susceptibility to carbapenems was the least frequent among the tested antibiotic classes. No isolate resisted meropenem, while only one showed intermediate resistance to ertapenem and seven to imipenem. Again, no carbapenemase-encoding genes were found in the genome of any of the non-susceptible isolates. Interestingly, however, when using the epidemiological cutoff value of the inhibition zone (≤ 27 mm) set for meropenem in the European Union [22], 17 isolates (22%) were classified as non-wild type, which indicates some mechanism of carbapenem resistance could be emerging in the studied population.

In relation to other public *Salmonella* genomes from Mexico, clinical cases were also the main source of beta-lactam-resistant *Salmonella:* χ^2^=72.5, P<0.0001, OR=12.8, 95CI 4.5-36.5 (Fig 3). Again, bovines were the most important reservoir, among food related samples, of these AMR genotypes: χ^2^=39.3, P<0.0001, OR=6.1, 95CI 3.0-12.7. Class-A extended spectrum beta-lactamase (ESBL)-encoding genes (i. e. *bla*CARB, *bla*TEM) were the most abundant among isolates from all sources. Genes encoding class-C ESBLs (i. e. *blaCMY*) were also present in a small proportion of isolates from clinical cases (4.6%), vegetables (0.3%), and aquatic environments (1.6%), while the *blaOXA* gene, which encodes a class-D ESBL, was only detected in 9% of clinical strains. Fortunately, class-B ESBL-encoding genes were not detected in the genome of the studied public isolates.

### Fluoroquinolone resistance

The main resistance mechanism we detected was that encoded by plasmid-mediated quinolone resistance (PMQR) genes, such as *qnrB19* and *qnrA1*, involved in quinolone target protection (DNA gyrase). This genomic feature, which is known to confer low-level quinolone resistance, was strongly associated with the intermediate resistance against ciprofloxacin observed in our study (χ^2^=36.8, P<0.0001). As mentioned before, we did not find statistical association between point-mutations in the QRDR and the observed phenotypes.

Among public *Salmonella* genomes, those from cattle origin were the major source of PMQR genes (Fig 3), with 32.5% of these isolates carrying *qnr* alleles (χ^2^=196.9, P<0.0001, OR=13.8, 95CI 8.5-22.2). Conversely, the proportion of PMQR-positive isolates was 18.2% in clinical strains, while it ranged from 1.6 to 6.0% across isolates from vegetables, seafood, and aquatic environments. Public genomes encoded additional quinolone resistance factors, such as efflux mechanisms (*oqxAB* genes). However, these genes were not as abundant as *qnr* alleles and were restricted mostly to clinical strains (13.6%) and a very small proportion of vegetable isolates (0.4%).

### Chloramphenicol resistance

Resistance to chloramphenicol in experimental isolates was mainly associated with efflux mechanisms (*floR* gene). This gene was present in most isolates of serovar Typhimurium (8/10), accounting for over 50% of chloramphenicol non-susceptible phenotypes. We detected a second resistance mechanism (antibiotic inactivation), encoded by a chloramphenicol acetyltransferase gene (*catA2*). However, this gene was present just in one isolate of serovar Give. Moreover, there were six non-susceptible isolates that were predicted as genotypically susceptible. Again, *ramR* mutations were strongly associated with chloramphenicol resistance (χ^2^=8.1, P=0.0045), OR=21.1 95CI 1.2-368.4.

As observed for previous antibiotic classes, there was a strong association between source and the proportion of phenicol-resistant *Salmonella* genotypes (χ^2^=187.7, P<0.0001) among public NTS isolates. Clinical strains were more likely to carry phenicols resistance genes than isolates from any other source: OR=11.7, 95CI 4.6-29.7. Likewise, cattle were the dominant reservoir, among food related samples, of these AMR genotypes (OR=15.6, 95CI 8.9-27.1). Still, the genomic AMR profile was similar across sources (Fig 3), with a predominance of efflux factors (i. e. *floR*) over enzymatic inactivation mechanisms (i. e. *catAB*).

### Folate pathway inhibitors resistance

Regarding folate pathway inhibitors, the most abundant resistance mechanism among experimental isolates was that encoded by *sul* alleles (i. e. *sul1* and *sul2*), which were present in 11 isolates. Conversely, just one isolate carried the *dfrA12* gene along with the *sul1* gene. Overall, there was a discrete proportion of experimental isolates that resisted trimethoprim-sulfamethoxazole (16.9%).

The resistance profile of public genomes again revealed isolates of bovine origin are more likely to carry trimethoprim-sulfonamide resistance genes as compared to those from vegetables, seafood and aquatic environments (χ^2^=142.0, P<0.0001), OR=11.7 95CI 7.1-19.2 (Fig 3). In all these sources, *sul* and *dfrA* alleles were the most common. In general, however, clinical isolates represent a more significant source of trimethoprim-sulfonamide resistant *Salmonella* as compared to those from food related isolates (χ^2^=175.9, P<0.0001), OR=11.5 95CI 4.7-28.2.

### Tetracycline resistance

Among experimental isolates, resistance to tetracycline was mainly associated with the presence of genes encoding efflux mechanisms (i. e. *tetABCG*). There were seven non-susceptible isolates that did not carry any of the known tetracycline resistance genes. However, all these isolates carried *ramR* mutations, which are associated with MDR profiles involving tetracyclines and other antibiotic classes [36].

Comparison with public *Salmonella* genomes revealed the highest proportion of tetracycline-resistance genotypes corresponded to bovine sources (χ^2^=189.1, P<0.0001), OR=10.9 95CI 7.1-16.9, as compared to vegetables, seafood, aquatic environments and even clinical cases (Fig 3). Regarding the genomic AMR profile, isolates from all sources carried some variants of the same efflux-mediated resistance determinants (*tet* alleles).

### MDR profiles

Among experimental isolates, MDR profiles were associated with the presence of a resistance island and, to a lesser extent, the occurrence of mutations in their genomes. For instance, most serovar Typhimurium isolates (9/10) carried SGI1. This genomic island harbors a class-1 integron containing multiple gene cassettes (i. e. *aadA2, blaCARB-2, floR, sul1, tetG*), which explains the observed MDR phenotypes of these isolates (Fig 4). This feature appears to be common in *S. enterica* ser. Typhimurium circulating in Mexico. The analysis of the whole set of Typhimurium isolates from Mexico deposited at NCBI (n=38) showed 55.3% of them have MDR genotypes. Among these, nearly 70% harbor multiple AMR genes associated with the ACSSuT phenotype. Refer to supplementary S3 Table for accession numbers and AMR genotypes of this group of isolates.

**Fig 4.**
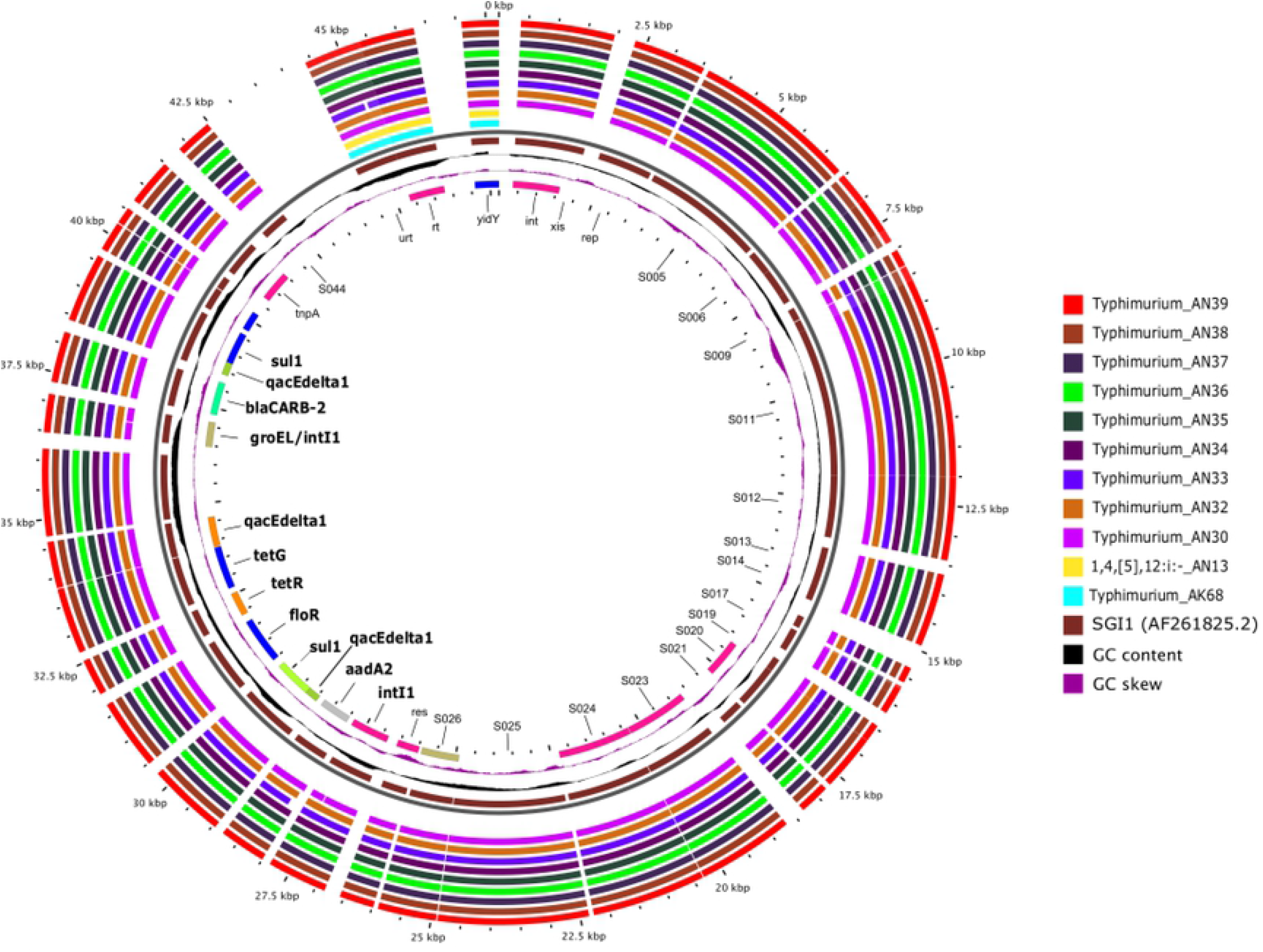
BLAST atlas analysis of SGI1 of 10 serovar Typhimurium isolates and one monophasic Typhimurium. The black slot corresponds to the backbone and the brown inner ring to the reference sequence. The most inner ring shows gene annotation, with AMR genes highlighted in bold text. The outer rings correspond to the serovars and sample names indicated. Regions with shared synteny between genomes and reference sequence are filled with intense color while blank spaces indicate lack of synteny. Refer to S1 Table for accession numbers and metadata of isolates.

As mentioned before, mutations in the *ramR* gene (Table 2) were strongly associated with MDR phenotypes as well. In fact, the probability of finding MDR strains was 50.5 times higher (95CI 2.9-881.3) in isolates carrying *ramR* mutations as compared to those lacking them.

Interestingly, plasmids had a rather discrete contribution to AMR phenotypes in general (Table 3). Only one third of the studied isolates (n=26) were predicted to carry seven different plasmids. Of these, three were small plasmids harboring few replication-related genes. The most abundant plasmid was the *Salmonella* virulence plasmid (pSLT), which was detected in all serovar Typhimurium isolates. The pSLT is a hybrid plasmid that carries a class-1 integron with several AMR gene cassettes against beta-lactams (*blaTEM*), chloramphenicol (*catA*), aminoglycosides (*aadA1, strAB*), and folate pathway inhibitors (*sul1*, *sul2*, *dfrA1*). However, the integron carried by our serovar Typhimurium isolates had a genomic context matching that of SGI1 instead of pSLT (see Fig 4 and supplementary S1 Figure).

**Table 3.**
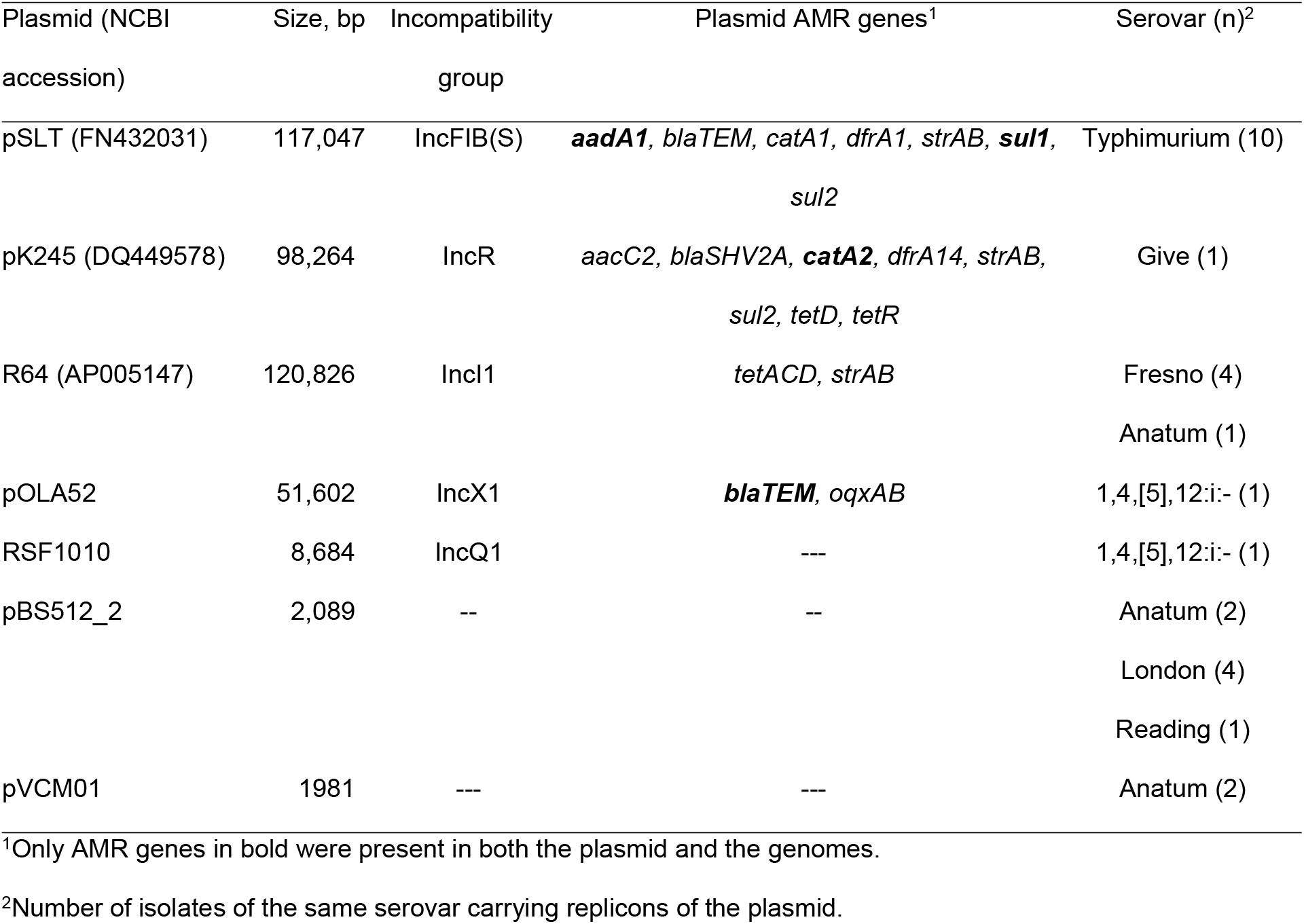
General features of the plasmids predicted per *Salmonella* serovar

Among the plasmids detected, the pK245 has the strongest resistance profile, carrying a class-1 integron with a sole resistance cassette (*dfrA14*), as well as another seven AMR genes (*tetAR, sul2, strAB, catA2, blaSHV-2*) distributed across the plasmid sequence. However, it is unlikely that pK245 plasmid was actually present in that isolate. Only 30% of this plasmid aligned to the genome of the single isolate (a serovar Give strain) carrying its replicons. Moreover, this isolate carried a chromosomal class-1 integron with seven resistance cassettes in a genomic context different from that of pK245 (see supplementary S2 Figure).

Replicons of the resistance plasmid R64 were also found in the genomes of serovar Fresno isolates (n=4), as well as one serovar Anatum isolate. This plasmid carries AMR genes against tetracyclines (*tetADR*) and aminoglycosides (*strAB*). Although over 70% of R64 aligned to the assembled genomes, none of its AMR genes were found in the genomes of isolates carrying its replicons, except *tetA*, which was detected in the serovar Anatum isolate (see supplementary S3 Figure). Finally, the monophasic Typhimurium isolate carried replicons of the pOLA52 plasmid. This is also a hybrid plasmid that harbors genes encoding a type IV secretion system, as well as AMR genes against quinolones (*oqxAB*) and beta-lactams (*blaTEM*). Although over 70% of this plasmid aligned to the referred genome, only the *blaTEM* gene was also present in the isolate, while *oqxAB* genes were missing (see supplementary S4 Figure).

Finally, the analysis of public genomes showed bovines and clinical cases are the most important sources, among those studied, of MDR NTS genotypes (Fig 3). In the other matrices, the proportion of MDR genotypes ranged from 2 to nearly 6%.

## Discussion

Our results confirmed previous observations of widespread resistance to older antibiotics (i. e. tetracyclines, penicillins) among NTS from different sources. For instance, in Mexico, previous studies from over 10 years ago, as well as recent research, report these phenotypes at very high frequencies (up to 90% or higher) in beef isolates [12, 15, 17, 37–39]. Likewise, these AMR phenotypes are also common in NTS isolated in developed countries, a phenomenon that is thought to be driven by the use of these antimicrobials in food-producing animals [40].

Regarding tetracyclines, selective pressure has likely resulted in the extensive acquisition of efflux mechanisms (*tetABCG* alleles), which are usually carried either in plasmids or in the chromosome of NTS [41]. Our results also support certain point mutations could sustain this phenotype in isolates lacking *tet* alleles, providing evidence of convergent evolution. As observed here, 100% of isolates that did not carry *tet* alleles had *ramR* mutations. These mutations are known to induce over expression of the AcrAB-RND efflux system [42], which confer resistance against multiple drugs, including tetracyclines.

Strikingly, results showed cattle seemed more relevant than any other source, including clinical cases, as a source of tetracycline resistance NTS. Although the number of clinical strains that were available for comparative analysis was low (n=22), these findings are relevant from a public health perspective. Tetracyclines are highly important antimicrobials [19] since they are among the few alternatives to treat human infections caused by *Brucella* spp., a pathogen that is associated with cattle.

The rate of resistance to penicillins observed here was not as high as that reported recently in NTS isolated from beef in Mexico [14, 15, 17, 43]. However, this variation is likely associated with the region where isolates originated. Hence, the relevance of cattle as a relevant source of penicillin resistant NTS could not be discarded, as shown by the genomic comparison with other public isolates from Mexico conducted here. In beta-lactam resistance, class-C and class-B ESBLs are the leading concerns. Class-C ESBLs confer resistance to most beta-lactams (except carbapenems) and are not inhibited by clavulanic acid, while class-B metallo-beta-lactamases confer the strongest resistance phenotype, involving all known beta-lactams and clavulanic acid [35]. Fortunately, resistance to 3GC, 4GC and carbapenems was rare among experimental isolates, which is in line with recent studies [9]. Apparently, food isolates in Mexico, as well as in the US and Europe [40], do not seem to be a significant source of NTS resistant to 3GC/4GC and carbapenems. Still, we observed some strains (10-15%) showed non-susceptibility to 3GC, 4GC and carbapenems, despite lacking ESBL-encoding genes. This could be the result of selective pressure and may lead to the emergence of resistance against these critically important antimicrobials, as shown by the moderate proportion of isolates (22%) exceeding the EUCAST’s ECOFF value for meropenem. In Mexico, 3GC used in human medicine (i. e. cefotaxime, ceftriaxone) are also approved, besides ceftiofur, to treat several bovine diseases [44]. Ceftiofur is associated with the emergence of ceftriaxone resistance among livestock and poultry NTS in the United States [40]. Hence, it is important to revise approval of both ceftiofur and other 3GC for veterinary use in Mexico. It has been reported that restrictions (voluntary or law-enforced) in 3GC use in the United States and Canada have helped reduce 3GC resistance among NTS associated with animals [40]. In any case, given the limited scientific evidence available in Mexico, continuous surveillance of NTS from foods is of upmost importance to support future risk management decisions.

We also observed decreased susceptibility to ciprofloxacin among experimental isolates, a phenomenon that has been associated with the presence of PMQR genes. Particularly, the *qnrB19* allele, which was present in over 50% of our experimental isolates, has also been reported as predominant among NTS strains from the United States [40]. Recent studies from Mexico have also documented high rates (36-44%) of decreased susceptibility to ciprofloxacin in beef NTS isolates [12], although these authors did not determine AMR genotypes. Moreover, a previous study by our research group documented *qnr* and *oqx* alleles were widely distributed among NTS isolated from cattle feces, carcasses and ground beef [17]. Furthermore, the genomic AMR profiling of public NTS isolates from Mexico conducted here, showed *qnr* alleles are widely disseminated among bovine isolates but not in those from vegetables, seafood, aquatic environments or even clinical strains.

In the context of Mexico’s cattle production, acquisition and conservation of PMQR genes in NTS is likely the result of selective pressure associated with the use of quinolones (i. e. enrofloxacin), which are approved to treat bovine respiratory, digestive, urinary and skin diseases, among others [44]. Since PMQR genes are carried in plasmids, this increases the risk of their dissemination under positive selective pressure. Many studies report quinolone resistance is rare in NTS isolated from cattle [3, 45]. Hence, it has been suggested that the use of these drugs in cattle could not be linked to the emergence of quinolone resistance among cattle NTS isolates [1]. However, PMQR genes are known to confer low-level quinolone resistance, which usually goes undetected when using CLSI breakpoints [46]. Therefore, we believe the role of PMQR genes as a contributing factor to quinolone resistance in NTS from cattle should not be minimized. Especially, considering PMQR genes have been associated with increasing resistance to quinolones in NTS isolated from foods [47, 48]. In summary, cattle appear to be a relevant source of low-level quinolone resistant NTS. However, to which extent this could result in the emergence of clinical fluoroquinolone resistance among cattle NTS isolates is yet to be determined.

Regarding aminoglycoside resistance, although the number of non-susceptible isolates in the experiment was low, the CLSI guidelines emphasize these antimicrobials may appear active *in vitro* but are not effective clinically against *Salmonella* and, thus, susceptible isolates should not be reported as such [21]. This may explain why isolates carrying *aadA* and *aph* alleles exhibited susceptible phenotypes, a phenomenon that has been observed in previous experiments [17]. Hence, it appears that WGS is a better tool to monitor aminoglycoside resistance as compared to AST. In that sense, our comparative genomics of public NTS from Mexico showed aminoglycoside resistant is less frequent in isolates from food related sources as compared to clinical cases. Resistance to chloramphenicol and trimethoprim-sulfamethoxazole was also frequently observed among experimental isolates (approximately 17 and 20%, respectively). These phenotypes were sustained by acquired AMR genes (i. e. *cat*, *flo, sul*, and *dfrA* alleles). In the case of chloramphenicol, *ramR* mutations, which are known to confer phenicol resistance [36], seemed to play a role as well.

These findings could be explained by the selective pressure exerted by the use of these drugs in food-producing animals. For instance, trimethoprim-sulfamethoxazole is approved as a wide spectrum antibiotic to treat all sorts of bacterial infections in livestock and poultry in Mexico [44]. Likewise, although chloramphenicol is no longer approved for these purposes, there are other phenicols (i. e. florfenicol) registered. This is likely the reason why phenicol resistance is consistently reported in NTS from animal foods in Mexico, in proportions that vary from moderate (16-23%) [12, 15, 39] to very high (>90%) [14, 43]. Our comparative genomics further supported this analysis since *floR*, which confer resistance to both phenicols [49], was the dominant resistance factor harbored by NTS from food related sources and clinical cases in Mexico.

Regarding MDR profiles, it is interesting to note the higher frequency of MDR isolates in ground beef as compared to lymph nodes. Perhaps this was influenced by the high number of MDR serovar Typhimurium isolates that were exclusively present in ground beef. However, some other factors could further explain these findings. For instance, research on ground beef consistently report high rates of MDR profiles in NTS, regardless of the relative serovar representation [12, 13, 15]. Conversely, studies involving NTS isolated from bovine lymph nodes usually report low levels of MDR phenotypes [8, 50]. To date, researchers have not been able to decipher the route of entry of NTS circulating in bovine lymph nodes. It has been suggested that fly bites could directly introduce the pathogen into the host’s blood [51]. Since this is reasonably likely to occur in feedlots, and NTS survives within the host cells [52], which are not affected by antibiotic treatments, these isolates may not be exposed to the same antimicrobial selective pressure as those circulating in the feces. Hence, they exhibit a weaker AMR profile. This study confirmed pan-susceptible phenotypes are more common among NTS isolates from bovine lymph nodes, while those from ground beef tend to have stronger AMR profiles, as well as higher rates of MDR phenotypes.

Among the genetic determinants associated with MDR phenotypes, SGI1 was the most frequently observed. It was present in all MDR serovar Typhimurium isolates (8/10), which accounted for 40% of the total number of MDR strains. The SGI1 contains a class-1 integron and multiple AMR gene cassettes (*aadA2, floR, tetG, bla-CARB-2, sul1*) conferring resistance to ampicillin, chloramphenicol, streptomycin, sulfonamide, and tetracycline [53]. This penta-resistance profile is typical of MDR *S. enterica* ser. Typhimurium DT104 carrying SGI1 [29] and was very similar to that observed here. Although SGI1 is not self-mobilizable, it can be transferred to other hosts by donor cells harboring conjugative plasmids [54]. In fact, SGI1 has been recognized as a key factor for the rapid dissemination of strains [55]. Unfortunately, although serovar Typhimurium is known to be dominant in NTS circulating in foods and clinical cases in Mexico, it has been poorly characterized in terms of its genomic AMR profile [9]. According to this review, the SGI1 associated ACSSuT phenotype has been reported in a modest proportion of MDR isolates (9.5%) collected from clinical cases, chicken, pork, and beef between 2006 and 2013. Analysis of public serovar Typhimurium isolates from Mexico deposited at NCBI (n=38) showed nearly 37% of them carry AMR genes supporting DT104-like or even stronger phenotypes. Nonetheless, further research is needed to establish if the high rate of DT104-like isolates observed here is a local dissemination phenomenon. Especially, considering half of our MDR Typhimurium isolates also showed decreased susceptibility to ciprofloxacin, which further amplifies the risks pose by these strains to human health. Moreover, it is important to better assess the contribution of point mutations and plasmids to AMR in NTS from foods, which appeared to be modest in the studied sample.

In summary, this research showed beef is a moderate source of MDR NTS harboring several AMR genes of human clinical significance. Among the sequenced public NTS isolates from foods and clinical cases in Mexico to date, those from cattle have the highest proportion of MDR genotypes. However, this should be interpreted with caution since the number of sequenced isolates from other species (i. e. chicken, pigs) and clinical cases is very limited in public databases.

Taken together, our results show it is vital to improve NTS control in apparently healthy animals to prevent its dissemination along the food chain and, consequently, human exposure to MDR strains. Particularly, we believe attaining significant improvements in AMR meat safety may require the identification and removal (or treatment) of product harboring MDR NTS, instead of screening for isolates showing resistance to individual antimicrobial classes. Such measures do not seem realistic, in the context of the meat industry of many countries, where testing practices are limited to *Salmonella* spp. isolation and confirmation. However, most nations have embraced the WHO global action plan on AMR. Thus, they should eventually set up existing technologies, such as WGS, which provide the shortest path to accomplish these goals.

## Acknowledgments

The authors appreciate the technical assistance and training on whole genome sequencing provided by the staff of the National Reference Center for Pesticides and Contaminants belonging to the General Directorate of Agri-Food, Aquaculture and Fisheries Safety of the national Service for Agri-food Health, Safety and Quality. We are also grateful for the support of laboratory technicians, as well as graduate and social service students from the Faculty of Veterinary Medicine, National Autonomous University of Mexico, in field sampling and laboratory analyses.

## Supporting information

S1 Table. NCBI accessions and metadata of our fully sequenced *Salmonella* isolates collected from bovine lymph nodes and ground beef (XLSX).

S2 Table. NCBI accessions, metadata and antimicrobial resistance genotypes of fully sequenced public *Salmonella* isolates from Mexico included in this study

S3 Table. NCBI accesions, metada and antimicrobial resistance genotypes of fully sequenced public *Salmonella enterica* ser. Typhimurium isolates from Mexico included in this study

S1 Figure. BLAST atlas of *Salmonella* virulence plasmid pSLT of ten experimental serovar Typhimurium isolates. The black slot corresponds to the backbone and the inner ring to the reference plasmid sequence and genes. Map generated with GView, version 1.7 through the tblastx program, with e-value=0.001, alignment cutoff=50, identity cutoff=70, and no filtering of low complexity sequences. Refer to S1 Table for isolates accessions and metadata.

S2 Figure. BLAST atlas of plasmid pK245 of one experimental serovar Give isolate. The black slot corresponds to the backbone and the inner ring to the reference plasmid sequence and genes. Map generated with GView, version 1.7 through the tblastx program, with e-value=0.001, alignment cutoff=50, identity cutoff=70, and no filtering of low complexity sequences. Refer to Table S1 for isolate accession and metadata.

S3 Figure. BLAST atlas of plasmid R64 of experimental serovar Fresno and Anatum isolates. The black slot corresponds to the backbone and the inner ring to the reference plasmid sequence and genes. Map generated with GView, version 1.7 through the tblastx program, with e-value=0.001, alignment cutoff=50, identity cutoff=70,and no filtering of low complexity sequences. Refer to S1 Table for isolate accessions and metadata.

S4 Figure. BLAST atlas of plasmid pOLA52 of one experimental serovar monophasic Typhimurium isolate. The black slot corresponds to the backbone and the inner ring to the reference plasmidsequence and genes. Map generated with GView, version 1.7 through the tblastx program, with e-value=0.001, alignment cutoff=50, identity cutoff=70, and no filtering of low complexity sequences.Refer to Table S1 for isolate accession and metadata.

## Author Contributions

### Conceptualization

María Salud Rubio Lozano, Orbelín Soberanis Ramos, Rubén Danilo Méndez Medina, Enrique Jesús Delgado Suárez.

### Data curation

Tania Palós Gutiérrez, Francisco Alejandro Ruíz López, Cindy Fabiola Hernández Pérez.

### Formal analysis

Tania Palós Gutiérrez, Cindy Fabiola Hernández Pérez, Nayarit Emérita Ballesteros Nova, Enrique Jesús Delgado Suárez.

### Funding acquisition

María Salud Rubio Lozano, Orbelín Soberanis Ramos, Marc W. Allard.

### Methodology

Cindy Fabiola Hernández Pérez, Francisco Alejandro Ruíz López, Enrique Jesús Delgado Suárez.

### Project administration

María Salud Rubio Lozano, Orbelín Soberanis Ramos.

### Resources

María Salud Rubio Lozano, Orbelín Soberanis Ramos, Francisco Alejandro Ruíz López.

### Software

Cindy Fabiola Hernández Pérez, Nayarit Emérita Ballesteros Nova, Enrique Jesús Delgado Suárez.

### Supervision

María Salud Rubio Lozano, Rubén Danilo Méndez Medina, Marc W. Allard, Enrique Jesús Delgado Suárez.

### Visualization

Tania Palós Gutiérrez, Enrique Jesús Delgado Suárez.

### Writing – original draft

Tania Palós Gutiérrez, Enrique Jesús Delgado Suárez.

### Writing – review & editing

Enrique Jesús Delgado-Suárez, Tania Palós-Guitérrez, Francisco Alejandro Ruíz-López, Cindy Fabiola Hernández Pérez, Nayarit Emérita Ballesteros-Nova, Orbelín Soberanis-Ramos, Rubén Danilo Méndez-Medina, Marc W. Allard, María Salud Rubio-Lozano.

